# Psilocybin differentially shifts whole-brain network dynamics during information processing in autism

**DOI:** 10.64898/2026.11.12.765347

**Authors:** Yan Ge, Tobias P. Whelan, Mihail Dimitrov, Claire L. Ellis, Francesca Moruzzi, Francesca M. Ponteduro, Johanna Kangas, Nermin Khalil, Fernando Zelaya, Glynis Ivin, Sunniva Fenn-Moltu, Nicolaas A. Puts, Declan G.M. Murphy, Eileen Daly, Grainne M. McAlonan, Dafnis Batalle

**Author notes:** Corresponding authors: Dr. Dafnis Batalle, Department of Forensic and Neurodevelopmental Science, Institute of Psychiatry, Psychology and Neuroscience, King’s College London 16, De Crespigny Park, SE5 8AF, London, UK, Professor Grainne McAlonan, Department of Forensic and Neurodevelopmental Science, Institute of Psychiatry, Psychology and Neuroscience, King’s College London 16, De Crespigny Park, SE5 8AF, London, UK. Joint senior authors (Grainne M. McAlonan, Dafnis Batalle).

## Abstract

Autistic people process information differently, yet conventional analyses of brain activity during information processing shows few differences. However, how brain-wide intrinsic connectivity networks (ICNs) are engaged during processing is rarely considered, nor are the dynamics of brain states during tasks and their modulation by drugs such as psilocybin (which alters the activity of large-scale brain networks). Here, during shape- and face-matching, we confirmed no autistic/non-autistic differences in conventional task-dependent activity, but observed reduced default mode and salience network strength in autistic participants. Following 5 mg oral psilocybin we observed a differential effect of psilocybin between autistic and non-autistic participants characterised by a lower salience network strength in non-autistic participants but not in autistic participants. Our main analysis of ICN dynamics identified four transient brain states: face-processing, global-hypoactivity, salience-attention, and default-mode-executive networks. These states occurred in both groups under placebo and psilocybin conditions. However, in autistic participants, psilocybin shifted whole-brain dynamics away from salience-attention networks towards self-referential and task-execution networks, particularly during shape-matching. These findings demonstrate that psilocybin differentially modulates large-scale brain dynamics in autistic and non-autistic adults and provide a rationale to assess whether shifting brain dynamics during information processing with psilocybin has potential clinical benefits in autism.

## Introduction

Autism has been conceptualised as an information processing condition^1^. Whole-brain intrinsic connectivity networks (ICNs) support fundamental cognitive processes involved in information processing^2^ and have been extensively characterised in autism during rest^3–7;^ however, little is known about how these large-scale networks reconfigure during active information processing. This distinction is important because resting-state organisation may not capture the network responses elicited by cognitive or sensory demands.

Most studies have focused on time-averaged (static) functional connectivity; however, brain functional activity is not static and fluctuates over time. Brain activity can be conceptualised as transitions through transient states of dynamic connectivity, providing a framework to investigate information processing beyond static connectivity. The few studies that have characterised atypical brain dynamics in autism during rest, typically report a reduced dynamic range of transient states with fewer transitions and atypical dynamic characteristics compared to non-autistic controls^8–11^. However, it remains unclear whether autistic brain dynamics are altered while processing information in standardised tasks.

There is also limited understanding about how neurotransmitter systems modulate task-related brain dynamic activity in autism. This question is especially important, given evidence suggesting that differences in the autistic brain may be more apparent in responses to neurochemical perturbation^12–14^ than in baseline neural function^15,16^. Previous findings suggest that the 5-HT_2A_ agonist psilocybin alters brain network connectivity differently in autistic participants^14^, and that it modulates brain network dynamics in (assumed) non-autistic people^17–22^. Comparing psilocybin-induced changes in autistic and non-autistic individuals can therefore clarify how serotonergic signalling shapes dynamic network reconfiguration during information processing in autism.

Here, we used a shape- and face-matching task^23^ to examine the static and dynamic reconfiguration of brain networks during non-social and social information processing in autistic and non-autistic adults. We first assessed conventional task-evoked BOLD responses associated with face-processing in autistic and non-autistic adults **(Fig. 1A)**. Then we characterised ICN spatial maps and core network strength of each ICN during the whole task **(Fig. 1B-C)**. Next, we characterised whole-brain state transitions of ICN co-activation across the task and examined baseline group differences **(Fig. 1D)**. We then tested the hypothesis that psilocybin would differentially alter brain network engagement and brain-state dynamics during shape- and face-processing in autistic and non-autistic adults **(Fig. 1E)**.

**Figure 1.**
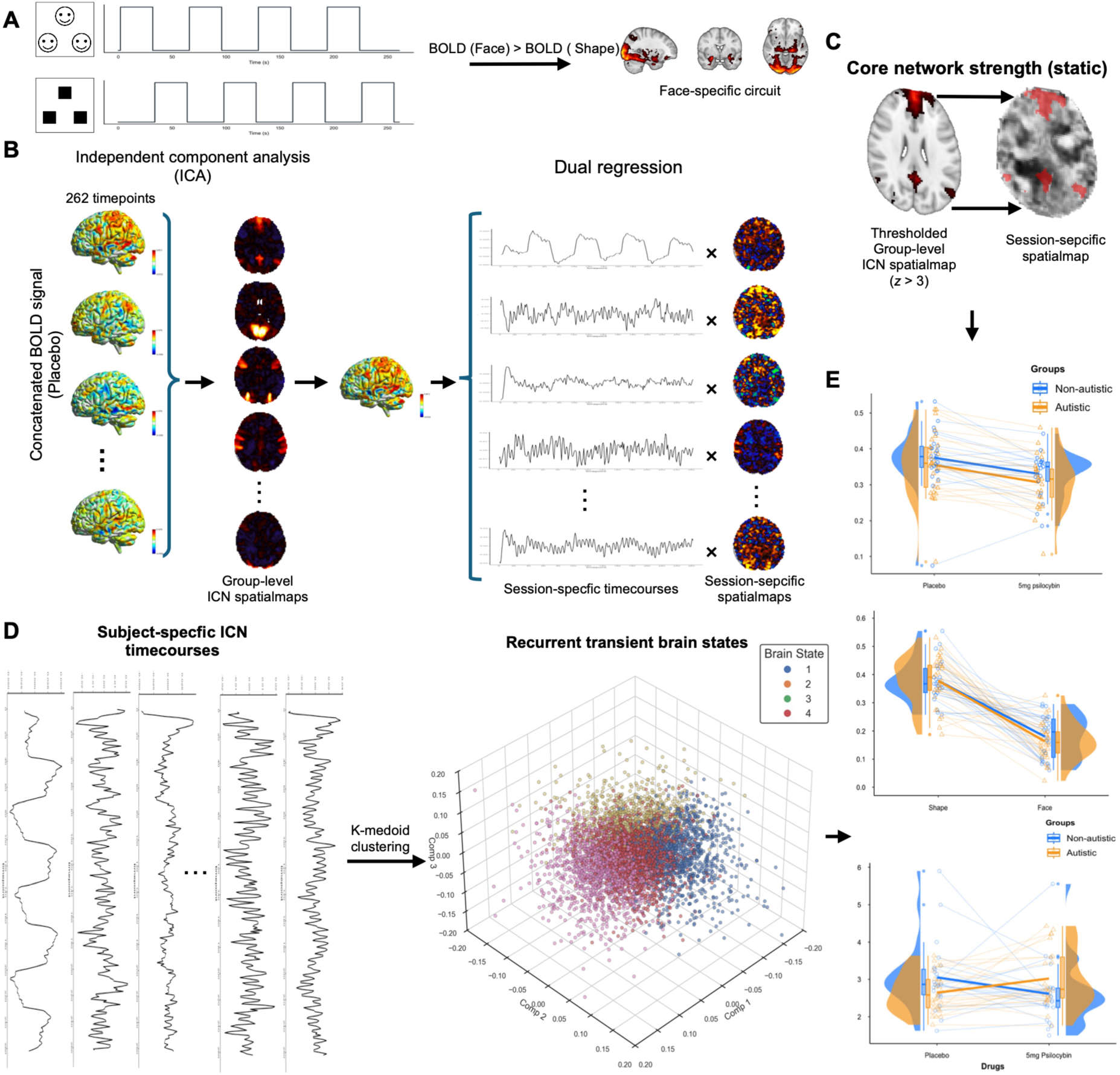
Overview of the methodological pipeline. **A)** BOLD response specific to face-processing was compared between autistic and non-autistic individuals and between conditions (placebo and 5 mg psilocybin). The facial images shown in this figure are illustrative computer-generated examples. During the actual task, participants viewed real emotional face stimuli^24^. **B)** Independent component analysis (ICA) was employed to extract group-level spatial maps of intrinsic connectivity networks (ICNs) from fMRI data acquired during the shape- and face-matching task after participants taking placebo. Session-specific timecourses and spatialmaps of ICN activity were obtained via dual regression from all fMRI sessions (after both placebo and 5 mg psilocybin). **C)** Core network strength examined average contribution strength of voxels within a thresholded ICN spatialmaps to this ICN (*z* > 3). **D)** Based on session-specific ICN amplitude timecourses, every timepoint was clustered into 4 recurrent transient brain states. We projected timepoints into the space of first three principal component (computed by principal component analysis (PCA)) (middle panel) for visualisation purposes. ICN dynamic features were characterised as fractional occupancy and dwell time. **E)** We compared core network strength and ICN dynamic features between autistic and non-autistic groups, between shape- and face-matching blocks, and between placebo and 5 mg psilocybin.

## Results

### Sample

In this study we analysed a sample of 33 autistic and 28 non-autistic participants (mean age (± SD) = 29.30 (± 8.54), female n = 29). fMRI data were acquired during performing a face-matching task^23^ after placebo or after a pharmacological challenge of low dose psilocybin (2 mg or 5 mg) on separate visit days, following a pseudo-randomised double-blind cross-over study design^25^ (see more details in the **Methods section**). The current analyses only included 5 mg psilocybin and placebo sessions as we previously reported non-significant effects of 2 mg psilocybin on functional connectivity^14^. After the exclusion of sessions with excessive head motion (5 sessions) and low behavioural performance (2 sessions), the final dataset comprised 102 sessions from 61 participants (autistic: n = 33): 56 sessions after placebo (autistic: n = 31) and 46 (autistic: n = 25) after 5 mg psilocybin. Autistic and non-autistic participants did not differ in age, task performance accuracy, reaction time, or head motion (**Table 1**). Across groups, 5 mg psilocybin significantly increased the mean reaction time (*t* = 2.36, *p* = 0.02) and subjective effects (*t* = 6.29, *p* < 0.001) measured with the 5-Dimensional Altered States of Consciousness Rating Scale (5D-ASC)^26^.

**Table 1.** Demographic information and behavioural performance of the Participants.

| Characteristics | Placebo |  |  |  | 5 mg psilocybin |  |  |  | Paired t test<sup>a</sup> |  |
|---|---|---|---|---|---|---|---|---|---|---|
|  | Mean (*SD*) / *n* (%) |  | t test / Chi-Square test |  | Mean (*SD*) / *n* (%) |  | t test / Chi-Square test |  |  |  |
|  | Non-autistic | Autistic | *t (*95% CI*) / χ<sup>2</sup>* | *p* | Non-autistic | Autistic | *t (*95% CI*)* | *p* | *t (*95% CI*) /χ<sup>2</sup>* | *p* |
|  | *N* = 25 | *N* = 31 |  |  | *N* = 21 | *N* = 25 |  |  |  |  |
| Age (years) | 28.80 (8.45) | 29.94 (9.04) | -0.48<br>(-5.84 – 3.57) | 0.63 | 29.14 (8.03) | 29.56 (9.21) | -0.16<br>(-5.54 – 4.70) | 0.87 |  |  |
| Gender |  |  |  |  |  |  |  |  |  |  |
| Female | 14 (56.00%) | 14 (45.16%) | 0.28 | 0.60 | 8 (38.10%) | 10 (40.00%) | 0 | 1.00 |  |  |
| Male | 11 (44.00%) | 17 (54.84%) |  |  | 13 (61.90%) | 15 (60.00%) |  |  |  |  |
| Accuracy (%) | 97.00 (4.21) | 97.38 (3.99) | -0.34<br>(-2.60 – 1.84) | 0.73 | 97.32 (5.11) | 98.50 (2.04) | -0.99<br>(-3.62 – 1.27) | 0.33 | 0.98<br>(0.29 – 0.82) | 0.34 |
| Reaction time (seconds) | 1211.59 (276.73) | 1149.54 (380.01) | 0.71<br>(-114.16 – 238.26) | 0.48 | 1315.64 (260.20) | 1297.79 (291.28) | 0.22<br>(-146.12 – 181.82) | 0.83 | 2.36<br>(17.74 – 228.60) | **0.02\*** |
| 5D-ASC total score | 36.24 (49.22) | 39.45 (30.70) | -0.28<br>(-26.04 – 19.62) | 0.78 | 149.29 (112.92) | 149.72 (115.34) | -0.01<br>(-68.51 – 67.64) | 0.99 | 6.29<br>(78.26 – 152.42) | **< 0.001\*\*** |
| Mean FD | 0.12 (0.04) | 0.13 (0.04) | -0.65<br>(-0.03 – 0.01) | 0.52 | 0.12 (0.04) | 0.12 (0.04) | -0.97<br>(-0.02 – 0.01) | 0.33 | 1.60<br>(-0.00 – 0.02) | 0.12 |

**Notes.** a. Paired t-tests were performed to compare task performance and 5D-ASC scores under placebo and 5 mg psilocybin. *SD*: standard deviation. 5D-ASC: 5-Dimensional Altered States of Consciousness Rating Scale. *CI*: confidence interval of mean difference. FD: framewise displacement. \* *p* < 0.05, \*\* *p* < 0.01

### Distinct recruitment of the salience and default mode network at baseline

Prior to analysis on ICNs, we performed conventional analysis on BOLD response to face-matching (**Fig. 2A** and **Supplementary Methods**). We did not find significant baseline group difference in BOLD response to faces nor drug effects of psilocybin on BOLD response in either group (see **Supplementary Results** for further details).

**Figure 2.**
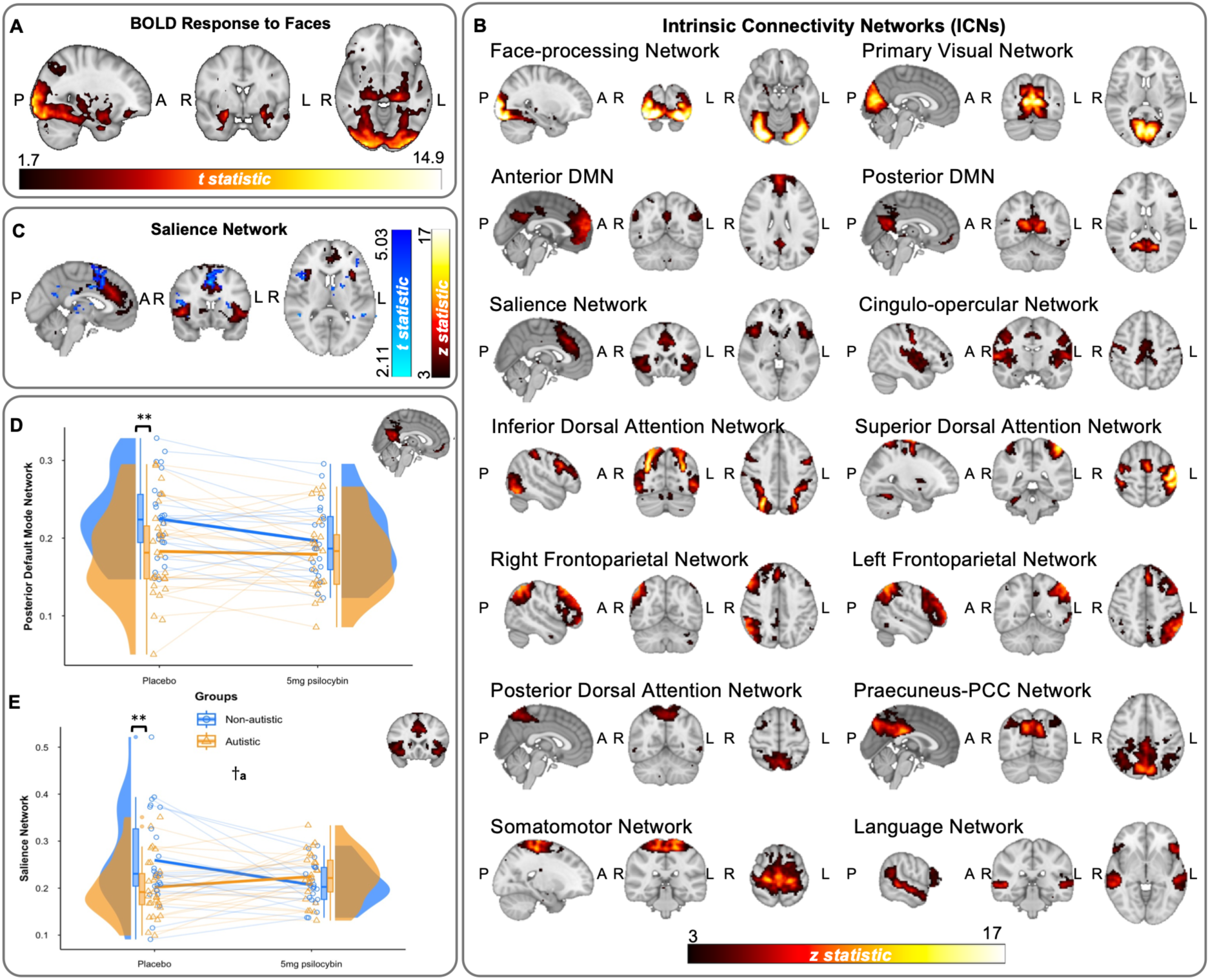
Static functional connectivity features. **A)** Voxels showing significantly stronger BOLD response to face-matching compared to shape-matching across all participants (one-tailed *p*_FWE_< .025). **B)** Intrinsic Connectivity Networks (ICNs) obtained with group ICA from fMRI data acquired during a shape- and face-matching task under placebo condition including both autistic and non-autistic participants. z-statistics were thresholded at *z* > 3. L: left. R: right. P: posterior. A: anterior. DMN: the default mode network. PCC: the posterior cingulate cortex. **C)** Voxels (in blue) showing significantly weaker contribution to the salience network (red) in autistic participants compared to non-autistic participants (contrast: non-autistic > autistic, one-tailed *p*_FWE_< .025). L: left. R: right. P: posterior. A: anterior. **D-E)** The core network strength of the posterior DMN and the salience network of autistic and non-autistic participants under placebo and 5 mg psilocybin conditions. ** indicates a significant group effect (*p* < 0.01). a. a significant group × drug interaction effect. † *p_FDR_* < 0.05. Solid lines represent changes from the mean under placebo to the mean under 5 mg psilocybin. The box bounds the interquartile range. Solid lines in the box indicate medians.

To investigate intrinsic connectivity networks supporting information processing, we extracted 14 ICNs with concatenated independent component analysis (ICA) ^27^(**Fig. 1B**). ICNs extracted included the face-processing network, the anterior and the posterior default mode network (DMN), the bilateral frontoparietal networks, and the salience network, among others (**Fig. 2B**). We obtained session-specific ICN amplitude timeseries (from dual regression stage 1 ^28^), i.e. activity amplitude of each ICN at each timepoint, and session-specific ICN spatial maps (from dual regression stage 2), i.e., voxel-wise strength of each ICN **(Fig. 1B)**. We first examined voxel-wise effects of group on ICN spatial maps at baseline. Autistic participants had weaker involvement of multiple regions belonging to the salience network (SN) than non-autistic participants at baseline (*p*_FWE_ < 0.025), including in superior frontal gyrus paracingulate gyrus, cingulate gyrus, frontal pole, and insular cortex (**Fig. 2C**).

We computed core network strength as the overall strength across voxels for each ICN per participant (**Fig. 1C**, see **Methods** for details). We used multiple linear regression to examine baseline group differences in ICN core network strength. At baseline, autistic individuals exhibited lower core network strength of posterior DMN (posterior DMN, *β* = -0.640, *t* = -2.800, *p* = 0.007, *p*_FDR_ = 0.050, **Fig. 2D**) and the salience network (*β* = -0.668, *t* = -2.937, *p* = 0.005, *p*_FDR_ = 0.050, **Fig. 2E**) than non-autistic individuals. Both effects survived FDR correction, but lay at the threshold of significance when adjusted for multiple comparisons of 14 ICNs. To address the heterogeneity within autism, we explored whether any observed differences in ICNs and their response to psilocybin were associated with dimensional measurements of autistic traits and co-occurring mental health symptoms (**Supplementary Table S1**), observing core network strength of the posterior DMN was correlated with a compound metric of autistic traits and mental health symptoms (**Supplementary Fig. S1** and **Supplementary Results**).

### Low dose psilocybin shifts ICN core network strengths

Across the whole scan, psilocybin reduced core network strength of multiple ICNs for both autistic and non-autistic participants, including the face-positive network and dorsal attention networks (see **Supplementary Results** and **Supplementary Fig. S2** for further details). A significant group × drug interaction effect was observed for salience core network strength (*β* = 0.905, *t* = 3.115, *p* = 0.003, *p*_FDR_ = 0.044, **Fig. 2E**). Post-hoc analyses suggested that 5 mg psilocybin reduced salience core network strength only in non-autistic participants (*β* = -0.456, *t* = -2.184, *p* = 0.044), although this effect did not survive FDR correction (*p*_FDR_ = 0.088). There was no significant voxel-wise drug or group × drug interaction effects on the spatial patterns of any ICN.

### Low dose psilocybin shifts whole-brain configuration patterns in autistic and non-autistic individuals

To examine whole-brain configuration dynamics, each timepoint was clustered into 4 recurrent transient brain states based on ICN amplitudes (i.e. ICN amplitude time courses from dual regression stage 1) with *k*-medoid clustering (**Fig. 1D**). The typical ICN activity of these 4 brain states (as represented by the 4 state medoids, **Fig. 3A**), can be summarised into a globally-hypoactivity state (State 1), a salience-attention state (State 2), a DMN-executive state (State 3), and a face-processing state (State 4) (see **Supplementary Fig. S3** for ICN activity levels in each state). The time courses depicting the percentage of participants in each brain state at baseline showed clear transition of brain dynamics between shape and face matching blocks (**Fig. 3A**), characterised by a very dominant face-processing state during face-matching blocks and a much more frequent recurrence of DMN-executive state and salience-attention state during shape-matching blocks. A global-hypoactivity state appeared with regular frequency through the whole task, for both face- and shape-matching blocks (see **Supplementary Table S2** for the probability of occurrence of each state). These medoids consistently occurred across different clustering resolutions (*k* = 3, 5, 6, and 7) and bootstrapped subsamples (see **Supplementary Fig. S4** for more details). We next examined whether psilocybin changed the dynamic properties of each of these transient states, i.e., their fractional occupancy and dwell time, and whether the effect was different in autistic and non-autistic participants and during shape- and face-matching conditions (see **Supplementary Fig. S5** for fractional occupancy of all the four brain states under placebo and 5 mg psilocybin conditions).

**Figure 3.**
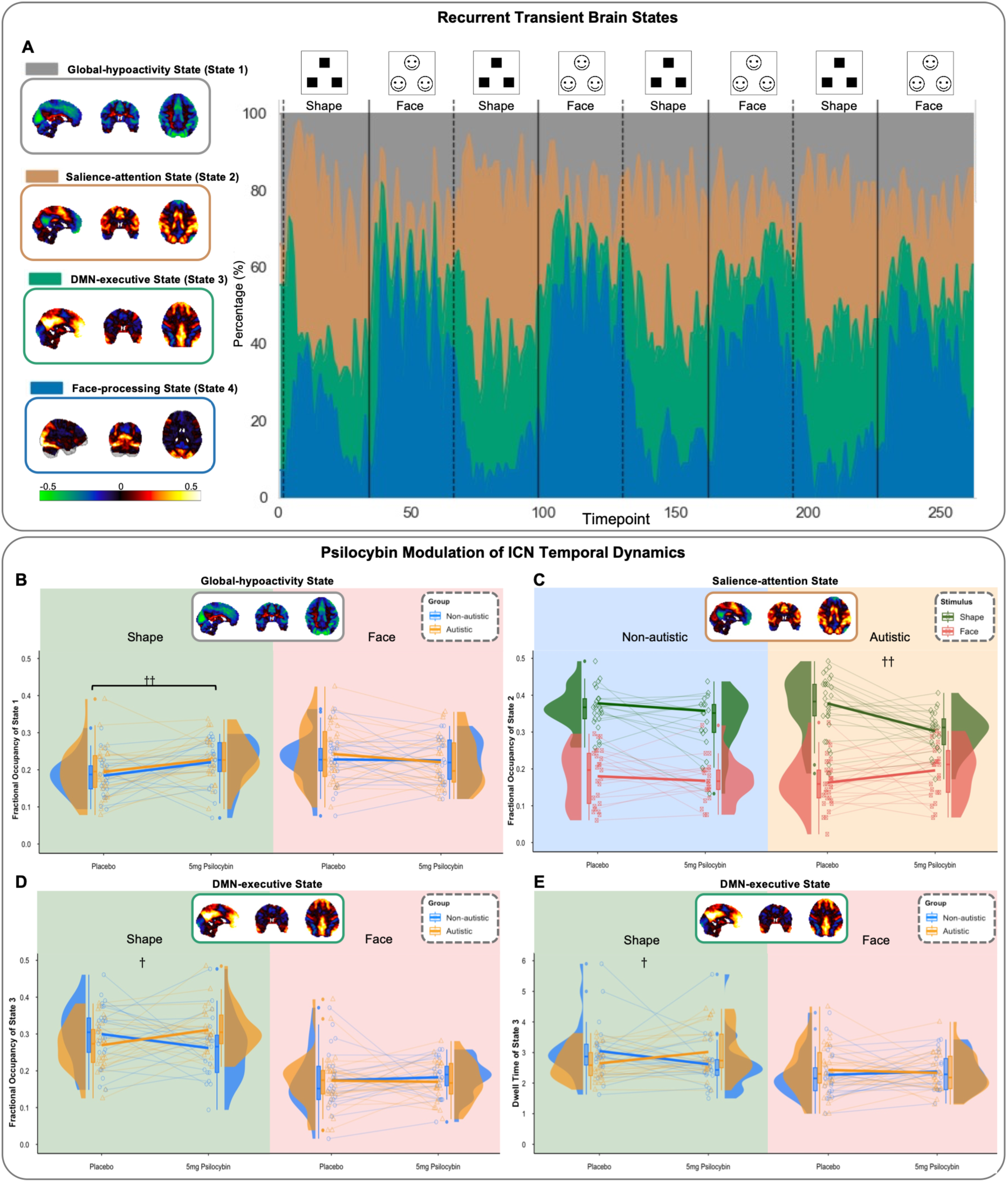
**A**) The Moment-to-moment activity of ICNs clustered into 4 recurrent brain states. The major spatial activity patterns of transient brain states, i.e., the medoids of each cluster, were shown on the left. The red-yellow colour bar indicates brain regions with increased activity, while the blue-green colour indicates brain regions with decreased activity. The time course of the percentage of participants in each brain state at baseline (placebo condition) was shown on the right. Dashed lines indicate the onsets of shape matching blocks, and solid lines indicate the onsets of face matching blocks. DMN: the default mode network. The facial images shown in this figure are illustrative computer-generated examples. During the actual task, participants viewed real emotional face stimuli^24^. **B)** The fractional occupancy of the global-hypoactivity state (State 1) under placebo and 5 mg psilocybin conditions in autistic and non-autistic participants. †† indicates a significant drug effect of psilocybin compared to placebo (*p_FDR_* < 0.01). Solid lines represent changes from the mean under placebo to the mean under 5 mg psilocybin. The box bounds the interquartile range. Solid lines in the box indicate medians. **C)** The fractional occupancy of the salience-attention state (State 2) under placebo and 5 mg psilocybin conditions in autistic and non-autistic participants. †† indicates a significant stimuli × drug interaction effect (*p_FDR_* < 0.01). Solid lines represent changes from the mean under placebo to the mean under 5 mg psilocybin. The box bounds the interquartile range. Solid lines in the box indicate medians. **D)** The fractional occupancy of the DMN-executive (State 3) state under placebo and 5 mg psilocybin conditions during shape and face blocks. † indicates a significant group × drug interaction effect (*p_FDR_* < 0.05). Solid lines represent changes from the mean under placebo to the mean under 5 mg psilocybin. The box bounds the interquartile range. Solid lines in the box indicate medians. **E)** The dwell time of the DMN-executive state (State 3) under placebo and 5 mg psilocybin conditions during shape and face blocks. † indicates a significant group × drug interaction effect (*p_FDR_* < 0.05). Solid lines represent changes from the mean under placebo to the mean under 5 mg psilocybin. The box bounds the interquartile range. Solid lines in the box indicate medians.

We found a significant drug × stimuli interaction effect on the fractional occupancy of the global-hypoactivity state (State 1) across the two groups (*β* = -0.711, *t* = -2.655, *p* = 0.009, *p*_FDR_ = 0.034)(**Fig. 3B**): 5 mg psilocybin significantly increased the fractional occupancy of the global-hypoactivity state during shape blocks (*β* = 0.559, *t* = 3.166, *p* = 0.003, *p*_FDR_ = 0.005) but not during face blocks (*β* = -0.212, *t* = -1.275, *p* = 0.208).

We observed a differential effect of psilocybin on autistic and non-autistic individuals in the fractional occupancy of the salience-attention state (State 2), modulated by the underlying stimuli (face vs shape matching), i.e., a group × drug × stimuli interaction (*β* = -0.848, *t* = -2.386, *p* = 0.018, *p*_FDR_ = 0.043, **Fig. 3C**). After post-hoc stratification analysis, we observed a significant drug × stimuli interaction effect on the salience-attention state (State 2) in autistic participants (*β* = 0.965, *t* = 4.096, *p* < 0.001, *p*_FDR_ < 0.001) whilst there was no significant drug effect in non-autistic participants. In autistic participants, 5 mg psilocybin decreased the fractional occupancy of the salience-attention state during shape blocks (*β* = -0.899, *t* = -4.110, *p* < 0.001, *p*_FDR_ < 0.001) but not during face blocks (*t* = 1.835, *p* = 0.079). In summary, psilocybin reduced occupancy of the salience-attention state (State 2) in autistic participants but not in non-autistic participants during shape matching.

There was also a significant group × drug × stimuli interaction effect on the fractional occupancy (*β* = -1.014, *t* = -2.312, *p* = 0.022, *p*_FDR_ = 0.043, **Fig. 3D**) and dwell time (*β* = -1.313, *t* = -2.660, *p* = 0.009, *p*_FDR_ = 0.035, **Fig. 3E**) of the DMN-executive state (State 3). After post-hoc stratification analysis, we observed a significant group × drug interaction effect on fractional occupancy (**Fig. 3D**) and dwell time (**Fig. 3E**) of the DMN-executive state during shape-matching blocks (*β* = 1.026, *t* = 2.919, *p* = 0.005, *p*_FDR_ = 0.022, and *β* = 1.006, *t* = 2.923, *p* = 0.005, *p*_FDR_ = 0.020, respectively) but not during face blocks. During shape blocks, psilocybin increased the fractional occupancy and the dwell time of the DMN-executive state in autistic (*β* = 0.584, *t* = 2.513, *p* = 0.018, *p*_FDR_ = 0.027 and *β* = 0.537, *t* = 2.551, *p* = 0.017, *p*_FDR_ = 0.033, respectively) but not in non-autistic participants (*t* = -1.550, *p* = 0.137, and *t* = -1.621, *p* = 0.118, respectively). The differential shifts (i.e., 5 mg psilocybin – placebo) in ICN dynamic features were associated with a compound metric of autistic traits and mental health symptoms (**Supplementary Fig. S6** and **Supplemenary Results**). Overall, these findings indicate that psilocybin shifts whole-brain dynamics in autistic participants away from salience-attention states towards DMN-executive states during non-social information processing.

## Discussion

Our main results showed that low dose psilocybin increased the occurrence of a deactivated brain state in both groups, and differentially altered whole-brain dynamic states in autistic and non-autistic adults (drug × group × stimulus interactions in DMN-executive and salience-attention states). Specifically, psilocybin shifted brain dynamics of autistic participants away from salience-attention states and towards DMN-executive states during shape-matching. The dynamic features of those two states were not significantly changed by psilocybin in non-autistic individuals.

Following a more traditional analysis pipeline to characterise static BOLD response to face-matching, we didn’t observe significant baseline group differences. These results align with previous findings that BOLD activation of face-processing circuits in response to face stimuli are similar in autistic and non-autistic adults, including in the fusiform gyri, amygdala, hippocampus, and occipital pole^16,29–32^. We also did not find differential response to psilocybin in task performance or BOLD response to faces between autistic and non-autistic participants. Consistent with previous literature, 5 mg psilocybin prolonged the reaction time of participants^33,34^. In contrast to studies applying higher dose of psilocybin, we didn’t find significant effects of psilocybin on BOLD activity with whole-brain comparisons^34,35^.

We further examined whether the configuration of each ICN during the whole task (on average) are distinct in autistic participants, and whether psilocybin differentially ‘shifted’ them. At baseline, we observed lower core network strength of the salience network and DMN in autistic adults, indicating reduced activation of those networks. The salience network is involved in detecting salient stimuli and guiding attention and cognitive processes^36,37^ and DMN supports emotional processing, mind wandering, and introspective thoughts^38^. Both networks have been broadly investigated during resting state in autism^39,40^ and have been found to be related to autistic traits^41–44^. Our findings are in line with under-recruitment of the salience network and DMN during tasks in autism^45,46^. We also found that reduced network strength of the DMN was associated with a compound metric of autistic traits and mental health symptoms, suggesting alterations in recruitment of the DMN might reflect dimensional changes in general autistic traits and co-occurring mental health symptoms. Importantly, differences emerged primarly at the level of large-scale organisation networks rather than within canonical face-processing circuits. Most differences were in brain association networks supporting salience detecting, attention switching, emotional processing, and self-referential thoughts, which have been suggested to underlie autistic differences in information processing^6^.

In line with previous studies that assessed resting state^18,47–49^ and face processing^50^, psilocybin consistently reduced the activation of DMN, dorsal attention network and face-processing circuits across autistic and non-autistic groups. These findings suggest that low dose psilocybin can desynchronise both task-specific and supporting networks during cognitive processing while most previous studies focused on high-dose psilocybin^51,52^. However, the core network strength of the salience network was only reduced in non-autistic participants, being shifted towards that of the autistic participants. Given the lower core network strength of the salience network in autism at baseline, this may reflect a ’baseline’ effect below which further lowering of core network strength by psilocybin in autism could not be detected. Alternatively, potential explanation for this is that the salience network consists of brain regions (i.e. the anterior cingulate cortex, anterior insular, bilateral caudate, and supplementary motor area) that are enriched in serotonin receptors targeted by psilocybin, including the 5-HT_2A_ receptor^53–55^. Lower 5-HT_2A_ receptor binding has been reported in these regions in autism^56^ and thus, the lack of salience network response to psilocybin in autistic participants may, at least in part, be explained by fundamental differences in their 5-HT system.

Going beyond static (time-averaged) features of single network configuration, we characterised four main transient brain states during the whole task. Despite baseline difference in within-network connectivity strength of the salience network, autistic participants showed similar whole-brain ICN coactivation patterns to non-autistic individuals during the whole task at baseline (placebo condition). Our results demonstrate that dynamic (temporal) features of ICN coactivation patterns were influenced by psilocybin particularly during shape-matching blocks. This may be due to the limited effects on BOLD response of the low dose of psilocybin we administered, potentially related to lower 5-HT_2A_ occupancy^57^, which was not able to shift task-specific brain configuration. Across both autistic and non-autistic participants, psilocybin significantly enhanced the occurrence of the global-hypoactivity state during shape blocks, i.e., increasing global ICN deactivation during non-social trials. This finding is in line with the effects of psilocybin we found on BOLD response to faces and with evidence that psilocybin reduces global brain activity, which has been lined to altered cognition^47^.

Despite no baseline difference, autistic brain dynamics were modulated differently by psilocybin during shape-matching blocks. Our time-resolved analysis demonstrated that during the shape-matching blocks, psilocybin shifted brain states in autistic participants from the occupancy of the salience-attention transient state, containing networks supporting attention directing salient and visual stimuli^58^, towards the DMN-executive transient state, containing networks supporting emotional processing, introspective thoughts, and task execution^54,59^. Similar to findings by Singleton et al^60^, the occurrence of transient brain states dominated by an active DMN were increased by psilocybin. Our findings suggest that, while processing non-social stimuli, autistic brain dynamics are more sensitive to modulation by low dose psilocybin. This may be partially explained by face blocks being dominated by a single face-processing state. Shape-matching blocks allow for a richer mix of dynamic brain activity, involving transitions among multiple large-scale states, creating greater opportunity to observe pharmacologically induced shifts in network configuration. The shifts in the occurrence of both the salience-attention state and DMN-executive state were both associated with autistic traits and co-occurring mental health symptoms. The findings reflect that altered 5-HT modulation of brain dynamics is related to dimensional changes in autistic behaviours and mental health, suggesting potential neurobiological mechanisms underlying altered whole-brain configuration in autism. As psilocybin has been shown to potentially benefit autistic individuals with anxiety and depression symptoms^61,62^, our findings may suggest potential clinical benefits of redirecting attentional resources in autism. Future studies are warranted to examine the relation between effects of psilocybin on brain dynamics, attention and autistic mental health.

There are several potential limitations in this study. Our sample only includes autistic adults with lower support needs, who showed no difference in the face- and shape-matching task performance compared to non-autistic groups. The lack of difference in activation of local brain circuits related to face-processing might be related to similar task performance^63^. Future studies may further investigate brain dynamics and response of autistic adults with higher support needs to neurochemical challenges, who might exhibit more diverse behavioural phenotypes. This would expand our understanding of the neurosignalling modulation and whole-brain organisation of autistic individuals, in a more heterogeneous cohort. Additionally, our study didn’t investigate the difference in serotonin systems explicitly, i.e. 5-HT receptors and transporters, between autistic and non-autistic individuals. We are not able to conclude the direct association between fundamental alterations in the 5-HT system and ICN engagement. Future studies may investigate whether differences in psilocybin targets, e.g. regional occupancy of 5-HT_2A_ receptors, are related to psilocybin-induced changes in within-network connectivity of the salience network and brain dynamics^64^ in autistic and non-autistic individuals. Moreover, the limited effect size of the association between behavioural phenotypes and psilocybin-induced changes in brain dynamics requires further studies with larger samples to validate this association.

## Conclusions

In summary, our key finding is that ICN dynamics in autistic adults during information processing respond to low dose psilocybin differently compared to non-autistic adults. In autistic adults, psilocybin shifted the occupancy from networks directing attention to salience and sensory stimuli towards networks supporting self-referential thoughts, emotional processing, and task execution while processing non-social information. Taken together, these findings suggest that serotonin targets of psilocybin modulate large-scale brain network dynamics responsible for attentional resource allocation, rather than face-specific responses in autism. This raises the possibility that there may be potential clinical uses of a psilocybin action on brain networks during information processing in autism which are worth exploring in future work.

## Methods

### Participants

We performed a secondary analysis of an fMRI dataset acquired during an adapted face matching task^23^ from a pseudo-randomised double-blind cross-over study^25^. The study recruited 30 (female n = 15) non-autistic and 37 (female n = 18) autistic adult individuals. Potential participants with major mental health conditions, genetic disorders associated with autism, intellectual disability, and/or any contra-indication for an MRI scan were excluded. All participants in the autism group had a clinical diagnosis of autism from a recognised assessment service. This work was part of a larger study^25^ that included a 2 mg psilocybin dose. Participants were asked to attend our centre three times and, on each visit, received 5 mg or 2 mg of COMP360 psilocybin, a proprietary synthetic formulation of crystalline psilocybin owned by Compass Pathways plc (London, United Kingdom), or (inactive) placebo in a pseudo-randomised order on separate visit days, at least 6 days apart to allow for drug wash out. The current study focused on 5 mg psilocybin and placebo sessions as non-significant effect of 2 mg psilocybin on functional connectivity and brain dynamics have been shown by previous research^14^.

### MRI Data Acquisition and Preprocessing

MRI data were acquired on a 3.0 Tesla MR Scanner (General Electric Premier). Structural MRI images were acquired with an 3D T1-weighted (T1w) Magnetization Prepared - RApid Gradient Echo (MP-RAGE) sequence (slice thickness = 1.00 mm, 208 slice locations, flip angle= 8°, field of view (FoV) = 256 × 256 mm, echo time (TE) = 2.96 ms, repetition time (TR) = 2658.90 ms, inversion time (TI) = 860 ms, matrix = 256 × 256). Task fMRI images were acquired using a gradient-echo echo planar imaging (EPI) sequence (slice thickness = 2.70 mm, slice spacing = 0 mm, 52 slices, flip angle = 60°, FOV = 221 × 221 mm, acquisition matrix = 82 × 82, TE = 32 ms, TR = 1000 ms, multiband factor 4, resolution = 2.7 × 2.7 × 2.7 mm^3^). Several pilot scans were carried out to arrive at an optimum multiband factor, which achieved a good balance between good temporal resolution (1s or less) and acceptable image quality. We collected one session of task fMRI for 270 s, comprising 270 time points on each visit. During the scan, participants performed an adapted face-matching task^23^ (see **Supplementary Methods** for a detailed description of the task). Two sessions with a poor face/shape-matching performance (accuracy < 75%) were excluded.

The functional MRI images were corrected for B0 distortion with the epi_b0_correct.py of AFNI 24.1.11^65^ and were then preprocessed with the standard pipeline of fMRIprep v23.0.1 (see **Supplementary Information** for full details of the pipeline). In brief, the task fMRI images were slice-time corrected using 3dTshift from AFNI^66^ and co-registered to the T1w-reference using bbregister of FreeSurfer^67^. The BOLD time-series were then resampled into the standard space, MNI152NLin2009cAsym. Framewise displacement (FD)^68^ was computed with Nipype v1.1.6^69^. After preprocessing with fmriprep, we smoothed the spatially-normalized preprocessed BOLD images with 4 mm full width at half maximum (FWHM) Gaussian kernel using FSL SUSAN^70^. Noise-regressors identified by automatic removal of motion artifacts using independent component analysis (ICA-AROMA)^71^, implemented in fMRIprep, were regressed out non-aggressively from the smoothed BOLD images with fsl_regfilt function in FSL v6.0.7.14 MELODIC (Multivariate Exploratory Linear Optimized Decomposition into Independent Components)^27^. Finally, nuisance signals - including the average time series of WM and CSF - were regressed out, and the data were high-pass filtered above 0.008 Hz with Nilearn v0.10.3^72^. Nine task fMRI sessions with a high level of motion mean FD > 0.3 mm or 20% FD > 0.3 mm were excluded.

### Behavioural questionnaires

Participants completed self-report measures of changes in states of waking consciousness induced by psilocybin or placebo, i.e. subjective effects, with 5-Dimensional Altered States of Consciousness Rating Scale (5D-ASC)^26^. They also completed self-report measures of autistic traits, including Adolescent/ Adult Sensory Profile (AASP)^73,74^, social and communicative behaviours with the Social Responsiveness Scale-2 (SRS-2)^75^ and Adult Repetitive Behaviours Questionnaire-2 (RBQ-2A)^76^, and reported mental health symptoms with Hospital Anxiety and Depression Score – depression subscale (HADS-D) and Hospital Anxiety and Depression Score – anxiety subscale (HADS-A)^77^. See **Supplementary Methods** for details.

### Data Analysis

#### Independent Component Analyses

We decomposed the fMRI data into voxel-wise components representing spatial patterns of coherent brain activity using group-level Independent Component Analysis (ICA)^78^. We extracted 25 components, representing a pragmatic balance between robustness and interpretability^79–81^, from the fMRI data across all participants under placebo condition. Among the 25 components, we identified 14 ICN components based on their spatial overlap with grey matter, their spatial patterns as biological patterns, activity time-series and power spectrum following the guidance of Griffanti et al.^82^. The spatial maps of those components were regressed into original fMRI voxel-wise BOLD timeseries under both placebo and psilocybin to retrieve the session-specific spatial maps of ICNs and session-specific timeseries of ICN activity at every timepoint via dual regression. We performed group ICA and dual regression with the MELODIC and the dual regression in FSL v6.0.7.14.

#### Dual Regression

Whole-brain voxel-wise general linear modelling with randomise permutation was performed with subject-specific spatial maps of ICNs to compare ICN patterns between autistic and non-autistic groups at baseline (placebo condition) using FSL v6.0.7.14 randomise^83^. The regression model consisted of main effect of group (including two contrasts of interest: autistic > non-autistic and autistic < non-autistic, using a voxel-wise FWE rate of one-tailed *p* < 0.025), age, sex, and mean FD (to control for residual motion effects) using FSL v6.0.7.14 randomise. To examine drug and group × drug interaction effect, we subtracted the subject-specific spatial maps under placebo from the spatial maps under psilocybin and then performed general linear modelling with randomise permutation. The regression model consisted of the overall effect of drug (including two contrasts of interest: average > 0 and average < 0, each using a voxel-wise FWE rate of one-tailed *p* < 0.025), main effect of group (including two contrasts of interest: autistic > non-autistic and autistic < non-autistic, using a voxel-wise FWE rate of one-tailed *p* < 0.025), age, sex, and mean FD using FSL v6.0.7.14 randomise.

#### Core Network Strength

The core network strength of each ICN were computed to measure the total within-network connectivity strength of ICNs, i.e., the average activation level of an ICN across voxels of each participant during the whole scanning period^79^. We obtained binary ICN masks by include voxels with z > 3 of the Z-transformed group-level ICN spatial maps^79,84^. Subsequently, the core network strength was calculated as a mean beta value across voxels of a session-specific spatial map within the binary ICN map with FSL v6.0.7.14. We first performed multiple linear regressions to examine the baseline (placebo condition) differences in core network strength between autistic and non-autistic participants. We then compared the effects of psilocybin on core network strength between non-autistic and autistic participants using linear mixed-effects models with group (autistic and non-autistic) × drug (placebo and psilocybin conditions) interaction as a fixed effect and participant ID and session order as random effects. *p*-values were corrected for multiple comparisons of core network strength of 14 ICNs using the Benjamini-Hochberg false discovery rate (FDR)^85^. Where significant interaction effects were found, we performed a stratification analysis (autistic and non-autistic groups) with linear mixed-effects models as post-hoc analysis. *p*-values of stratification analyses were corrected for two groups.

#### ICN dynamics

To examine ICN dynamics, we employed *k*-medoid clustering to define transient brain states, i.e., the recurrent patterns of ICN dynamic activity. K-medoid clustering is a partitioning method to identify clusters with minimal sums of dissimilarities between points in the data^86^. A medoid is an actual data point with the smallest total distances to other points in a cluster. We applied *k*-medoid clustering to the ICN activity matrix of 262 timepoints × 14 ICNs × 102 task-based fMRI sessions (including placebo and 5 mg psilocybin conditions) including all participants. We used the Manhattan distance to compute the dissimilarity matrices for clustering. To determine the number of clusters, we used the Davies–Bouldin index and the Calinski– Harabasz index to select *k* = 4 as the optimal number of partitions (**Supplementary Fig. S7**). To assess the robustness of the identified brain states, we examined correlations of those medoids with the medoids obtained at different clustering resolutions (*k* = 3, 5, 6, and 7) and across bootstrap subsamples. We generated 1,000 subsamples, each containing 80% of the full dataset, and applied *k*-medoids clustering to each subsample using *k* = 4. We then calculated Pearson correlations between the medoids derived from each subsample and those derived from the full dataset. Each subsample medoid was matched to the most highly correlated full-dataset medoid, and the distributions of the resulting correlation coefficients were plotted across the 1,000 subsamples.

#### Dynamic features

The dynamics of ICN activity were then characterised by fractional occupancy and dwell time of identified brain states. Fractional occupancy is the probability of a brain state occurring during an fMRI session. It was calculated as the percentage of time points clustered into a certain brain state among all time points. Dwell time is the average duration of a certain brain state during an fMRI session. It was calculated as the average number of continuous time points clustered into a certain brain state for each session. Scripts of R codes available in the *stateR* package^87^ were used to extract these features.

We first performed linear mixed-effects models to examine the baseline (placebo condition) differences in fractional occupancy, and dwell time between autistic and non-autistic participants with session orders as a random effect. We then compared the effects of psilocybin on fractional occupancy, and dwell time between non-autistic and autistic participants and between shape and face blocks using linear mixed-effects models with group × drug × stimuli (shape and face conditions) interaction as a fixed effect and session order and participant ID as random effects. *p*-values were corrected for multiple comparisons of fractional occupancy and dwell time of 4 brain states using the Benjamini-Hochberg false discovery rate (FDR)^85^. Where significant interaction effects were found, we performed stratification analyses with linear mixed-effects models as post-hoc analysis. *p*-values of stratification analyses were corrected for two groups and two stimuli (face and shape).

We controlled our analyses for sex, age, and motion (mean FD). All continuous variables were standardised before being entered into equations. R 4.5.1 was used to perform *k*-medoid clustering, multiple linear regressions, and linear mixed models as well as computing fractional occupancy and dwell time.

## Code and data availability

Data used in this study can be accessed by reasonable request and subject to data ownership agreements. The analysis code is available at: https://github.com/Kiragy/Psilocybin-whole-brain-network-dynamics-during-information-processing-in-autism-.git

