## Supplementary Information for "Psilocybin differentially shifts whole-brain network dynamics during information processing in autism"

### Supplementary Results

#### Face-processing BOLD response

We first examined group (autistic vs non-autistic) and drug (psilocybin vs placebo) effects of BOLD response to face-matching in task-specific regional circuits, following a standard task-based fMRI investigation workflow: subject-level and group-level voxel-wise whole-brain general linear modelling (GLM) with “faces > shape” contrast. On group average, the well-documented face-processing circuit, fusiform gyrus, temporal fusiform cortex, occipital pole, and bilateral amygdala and hippocampus, showed greater activation during face- than shape-matching (voxel-wise family-wise-error (FWE) rate of one-tailed $p < 0.025$) at baseline (**Fig. 2A**). No significant voxel-wise FWE-corrected effects of the group (autistic/non-autistic), drug (placebo/psilocybin), or group × drug interaction were found.

#### Distinct recruitment of the default mode network is associated with autistic traits at baseline

We examined the association between altered core network strength of the salience network and posterior DMN with autistic traits and co-occurring mental health symptoms at baseline. As no single metrics can capture the heterogeneity of autism in social and sensory responsiveness, and the co-occurrence of anxiety and depression in autism, we computed principal component analysis on standardised scores of the Adolescent/Adult Sensory Profile (AASP), Social Responsiveness Scale-2 (SRS-2), Adult Restricted and Repetitive Behaviours Questionnaire-2 (RBQ-2A), and the Hospital Anxiety and Depression Scale – Depression Subscale (HADS-D) and Anxiety Subscale (HADS-A) (see **Supplementary Table S1** for available questionnaire scores of participants) and obtained the first principal component (PC1) as a variable reflecting autistic traits and co-occurring mental health symptoms (see **Supplementary Fig. S1A** for the PCA loading plot)**(Supplementary Fig. S1B)**. The lower core network strength of the posterior DMN was correlated with higher (more autistic trait) PC1 scores ($r = -0.306$, $p = 0.029$, 95% CI [-0.562, -0.051]) (**Supplementary Fig. S1C**), but the salience network was not correlated with PC1 scores.

#### Low-dose psilocybin shifts ICN core network strengths

Across the whole scan, psilocybin reduced core network strength of multiple ICNs for both autistic and non-autistic participants, including the face-positive network ($\beta = -0.574$, $t = -5.623$, $p < 0.001$, $p_{FDR} < 0.001$, **Supplementary Fig. S2A**), inferior dorsal attention network (DAN) ($\beta = -0.534$, $t = -2.515$, $p = 0.017$, $p_{FDR} = 0.021$, **Supplementary Fig. S2B**), posterior dorsal attention network ($\beta = -0.586$, $t = -3.549$, $p = 0.002$, $p_{FDR} = 0.011$, **Supplementary Fig. S2C**), anterior DMN ($\beta$ = -0.559, $t$ = -2.571, $p$ = 0.012, $p_{FDR}$ = 0.021, **Supplementary Fig. S2D**), and language network ($\beta$ = -0.427, $t$ = -2.699, $p$ = 0.012, $p_{FDR}$ = 0.033, **Supplementary Fig. S2E**).

The shifts in the core network strength induced by psilocybin were not significantly associated with autistic traits and co-occurring mental health symptoms.

#### Distinct alterations in whole-brain configuration patterns are associated with autistic traits at baseline

We further examined the associations of differential shifts in ICN dynamic features (5 mg psilocybin - placebo) and PC1 scores of autistic traits and co-occurring mental health symptoms. PC1 scores were significantly associated with the shifts in fractional occupancy of both the salience-attention state ($r$ = -0.360, $p$ = 0.037, 95% CI [-0.704, -0.023]) and the DMN-executive state ($r$ = 0.363, $p$ = 0.045, 95% CI [0.009, 0.717]) (**Supplementary Fig. S6**).

**Supplementary Table S1.** *Questionnaire and PC1 scores of the Participants.*

| Questionnaires | Mean (*SD*)/ *n* (%) | | t-test | |
|---|---|---|---|---|
| | Non-autistic | Autistic | *t* (95% CI) | *p* |
| | *N* = 22 | *N* = 26 | | |
| HADS-A | 5.91 (3.34) | 11.65 (4.52) | -5.06 (-8.03 – -3.46) | < 0.001** |
| HADS-D | 1.86 (2.44) | 6.42 (4.20) | -4.69 (-6.52 – -2.59) | < 0.001** |
| SRS-2 | 54.36 (8.14) | 72.54 (7.09) | -8.17 (-22.66 – -13.69) | < 0.001** |
| RBQ-2A | 27.36 (6.59) | 37.08 (6.07) | -5.28 (-13.43 – -6.00) | < 0.001** |
| AASP | 143.77 (24.38) | 174.42 (19.08) | -4.79 (-43.60 – -17.70) | < 0.001** |
| PC1 | -1.52 (1.09) | 1.29 (1.36) | -7.93 (-3.52 – -2.09) | < 0.001** |

**Notes.** HADS-A: Hospital Anxiety and Depression – Anxiety Subscale. HADS-D: Hospital Anxiety and Depression – Depression Subscale. SRS-2: the Social Responsiveness Scale-2. RBQ-2A: the Adult Repetitive Behaviours Questionnaire-2. AASP : Adolescent/ Adult Sensory Profile questionnaire. PC1: first principal component derived from the 5 questionnaire scores showed here with principal component analysis.

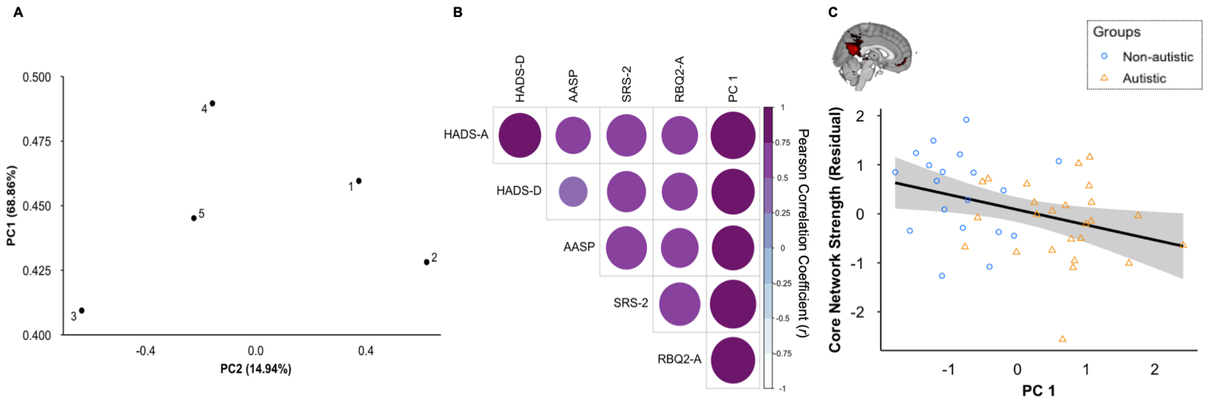


**Supplementary Fig. S1. A)** Loading plot of principal component analysis (PCA) on autistic traits and mental health symptoms. Variance proportions explained by each principal component are shown along each component axis. **B)** PC1 scores of baseline autistic traits and co-occurring mental health symptoms were highly correlated with each individual measure. **C)** The core network strength of the posterior DMN was significantly correlated with PC1 scores. We plotted the residuals of the core network strength after adjusting for age, gender, and mean FD in multiple linear regression models. The 95% confidence region is depicted by grey shading.

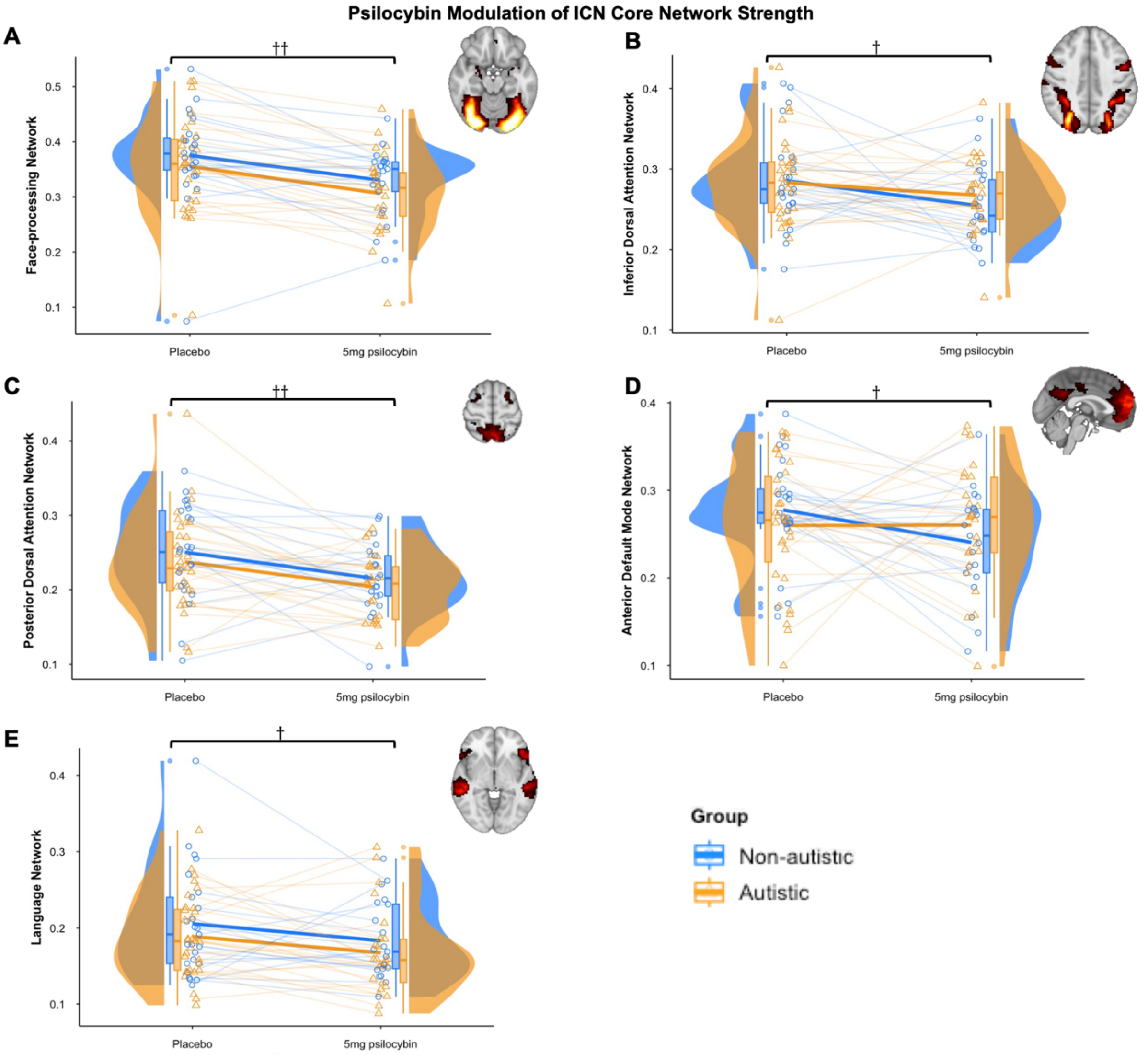


**Supplementary Fig. S2.** The core network strength of ICNs in autistic and non-autistic participants, under placebo and 5 mg psilocybin conditions. † $p_{FDR} < 0.05$. †† $p_{FDR} < 0.01$. Solid lines connecting placebo and 5 mg psilocybin conditions represent changes in the mean from placebo to 5 mg psilocybin. The box bounds the interquartile range. Solid lines in the box indicate medians.

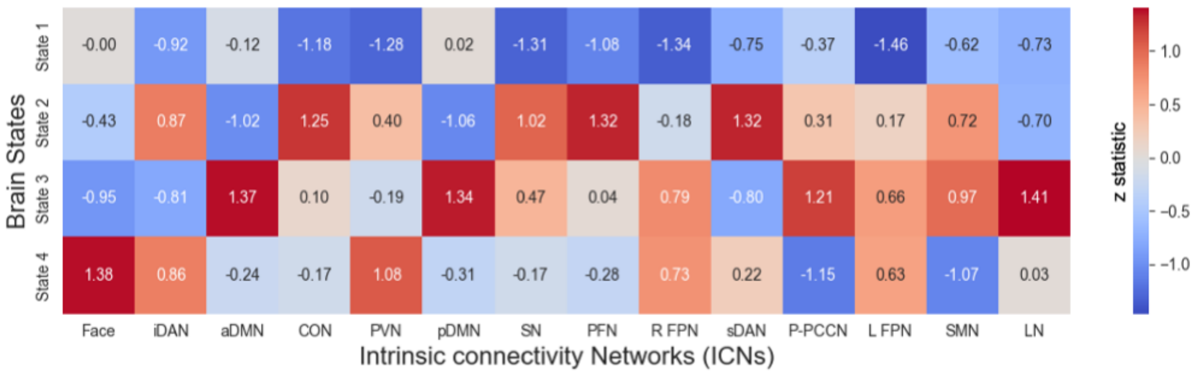


**Supplementary Fig. S3.** Heatmap of standardised ICN activity levels in each brain state. Face: face-processing network. The red-blue colour bar indicates the standardised level of ICN activity. iDAN: inferior dorsal attention network. aDMN: anterior default mode network. CON: cingulo-opercular network. PVN: primary visual network. pDMN: posterior default mode network. SN: salience network. PFN: parietal-frontal network. R FPN: right frontoparietal network. sDAN: superior dorsal attention network. P-PCCN: praecuneus-posterior-cingulate-cortex Network. L FPN: left frontoparietal network. SMN: somatomotor network. LN: language network.

**Supplementary Table S2.** *Probability of occurrence of each brain state during face- and shape-matching blocks after placebo.*

| | State 1 | State 2 | State 3 | State 4 |
|---|---|---|---|---|
| Shape | 18.99% | 37.81% | 28.31% | 14.90% |
| Face | 23.80% | 16.70% | 16.87% | 42.63 |

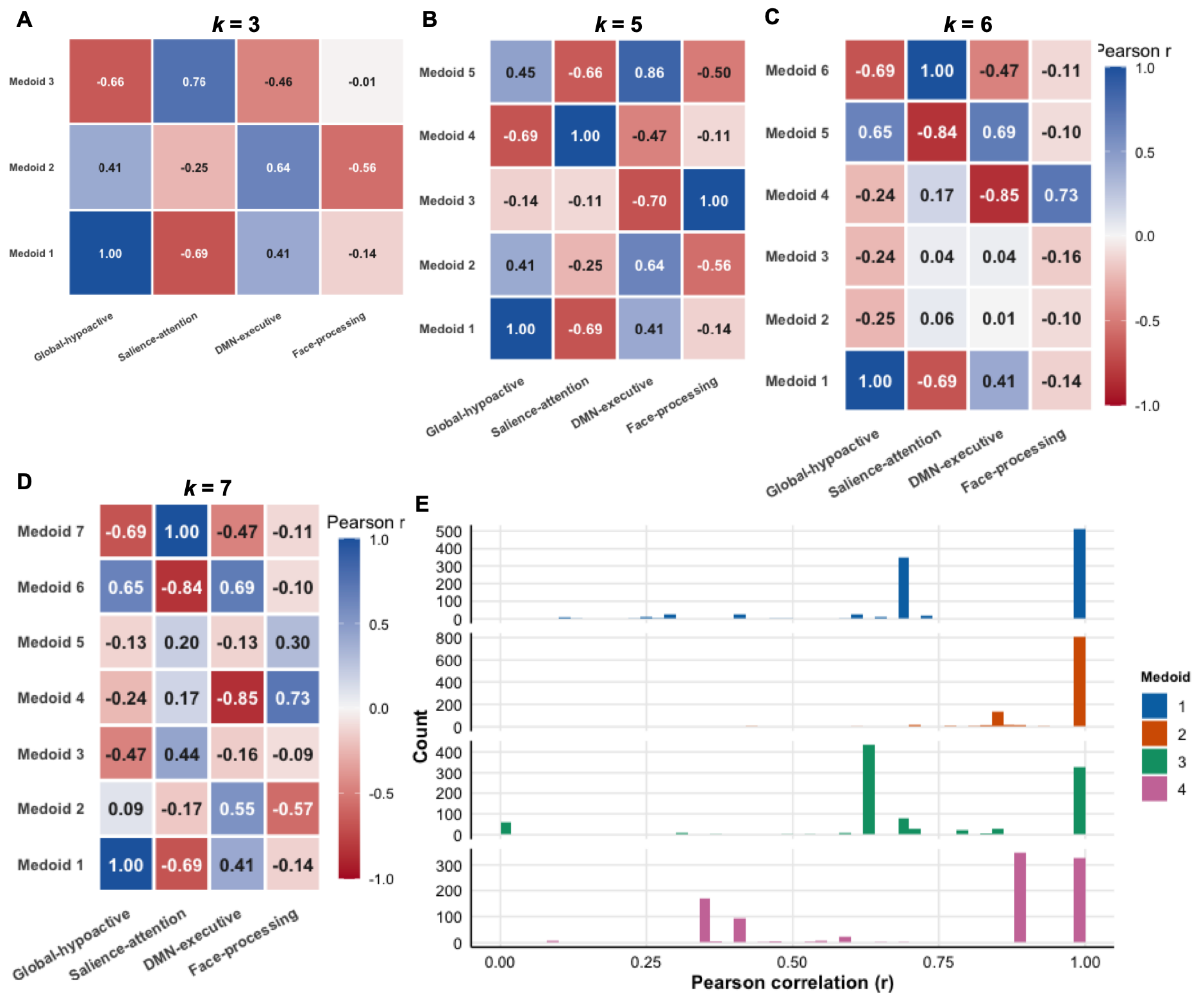


**Supplementary Fig. S4. A-D)** Pearson correlations between medoids obtained with *k* = 4 and those obtained at other clustering resolutions (*k* = 3, 5, 6, and 7). Medoids representing the same or highly similar brain states were consistently identified across clustering resolutions. **E)** Distribution of Pearson correlation coefficients between medoids obtained from 1,000 subsamples and those obtained from the full dataset, using *k* = 4. The medoids were consistently highly correlated across subsamples, demonstrating the robustness of the identified brain states.

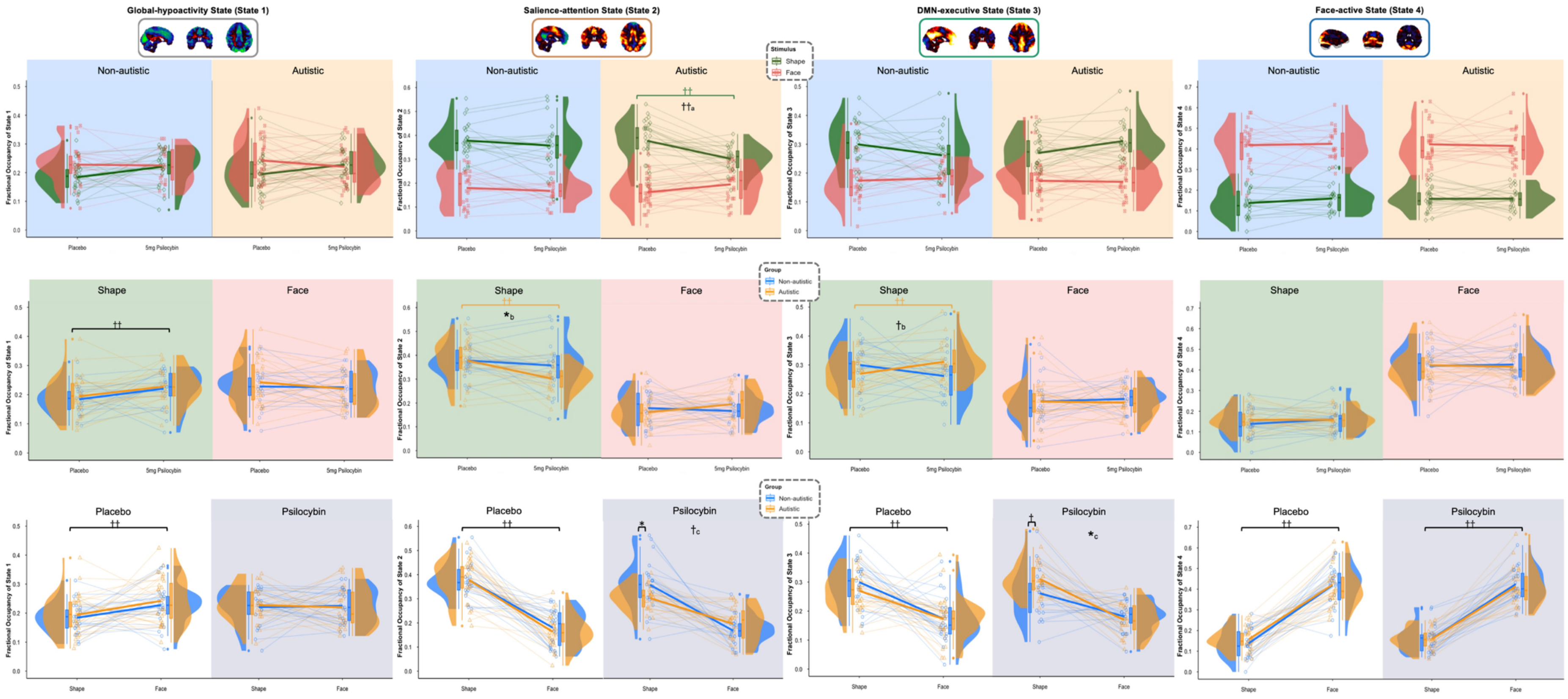


**Supplementary Fig. S5.** The fractional occupancy of the four transient brain states in autistic and non-autistic participants, during shape and face matching, and under placebo and 5 mg psilocybin conditions. * $p_{uncorrected} < 0.05$. † $p_{FDR} < 0.05$. †† $p_{FDR} < 0.01$. a. a stimuli × drug interaction effect. b. a group × drug interaction effect. c. a group × stimuli interaction effect. Solid lines connecting placebo and 5 mg psilocybin conditions/ shape- and face-matching blocks represent changes in the mean from placebo to 5 mg psilocybin/ from shape-matching to face-matching blocks. The box bounds the interquartile range. Solid lines in the box indicate medians.

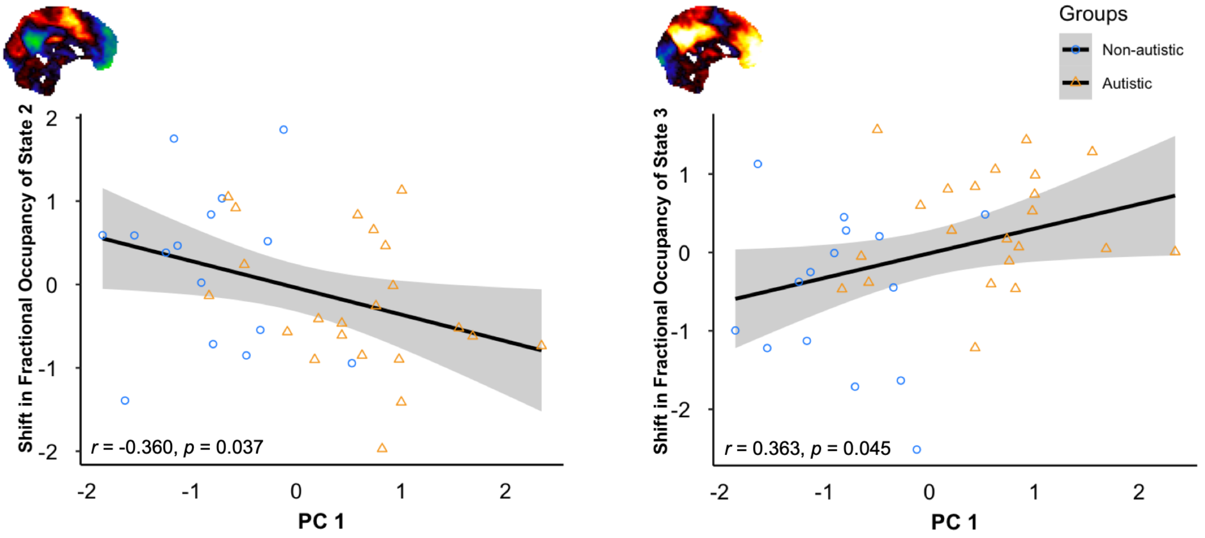


**Supplementary Fig. S6.** The association of psilocybin-induced shifts in the fractional occupancy of salience-attention states (State 2) and DMN-executive (State 3) with PC 1 scores of autistic traits and co-occurring mental health symptoms. We plotted the residuals of shifts in the fractional occupancy after adjusting for age, gender, and mean FD in multiple linear regression models. The 95% confidence region is depicted by grey shading.

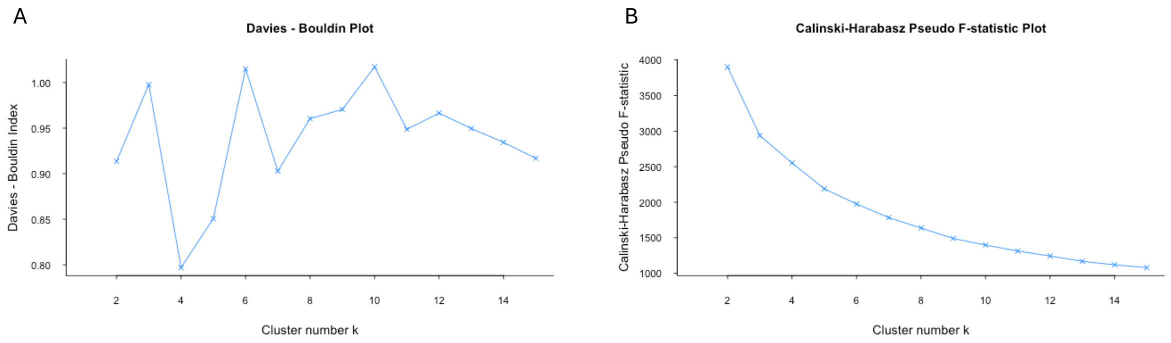


**Supplementary Fig. S7.** A) The Davies-Bouldin index corresponding to *k* from 1 to 15. B) The Calinski–Harabasz pseudo F-statistic corresponding to *k* from 1 to 15.

### Supplementary Methods

#### Face matching paradigm

On each trial, they were shown a target image (upper panel) and two comparison images (lower panel) and indicated the matching image via button press. Stimuli included sex-balanced emotional faces as experimental conditions and geometric shapes as controls. The face-matching and shape-matching blocks, each comprising six trials lasting 5 seconds, appear alternatively for four times in total.

#### Behavioural questionnaires

Changes in states of waking consciousness induced by psilocybin or placebo, i.e. subjective effects, were assessed with 94-item 5-Dimensional Altered States of Consciousness Rating Scale (5D-ASC)[1,2]. Participants reported their subjective effects approximately 4 hours (mean, 235 (± 25.4) mins) after the drug was administered. Participants rated the extent to which each item applies to their experience on a scale from 0 to 10. These scores are then summed to produce the 5D-ASC total score, with higher scores indicating stronger subjective effects induced by drugs.

Participants reported their behavioural response patterns to sensory stimuli with the Adolescent/ Adult Sensory Profile (AASP) questionnaire. It consists of four quadrants: low registration, sensation seeking, sensory sensitivity, and sensation avoiding[3], each including 15 items. Each item is rated with a 5-likert scale ranging from “almost never” to “almost always” and then summed up to produce the total AASP scores range from 60 to 300, with higher scores indicating more atypical sensory response patterns.

Social and communicative behaviours were self-reported by participants with the Social Responsiveness Scale-2 (SRS-2)[4]. The questionnaire includes five subscales: social awareness (8 items), social cognition (12 items), social communication (22 items), social motivation (11 items), and restricted and repetitive behaviours (12 items). Each item is rated with a 4-likert scale ranging from 0 to 3. The total scores are reported as T-scores ranging from 30 to 90. A higher score indicated more pronounced difficulties in reciprocal social behaviours.

Restricted and repetitive behaviours were self-reported by participants with the Adult Repetitive Behaviours Questionnaire-2 (adult RBQ-2)[5]. The questionnaire includes 20 items related to four factors, repetitive motor movements, adherence to routine, restricted interests, and unusual sensory interests. The items are 3-point likert, ranging from “Never or rarely” to “15 or more times daily” or “from Never or rarely through to “Marked or notable”. The scores of each item are summed up to produce total RBQ-2A scores ranging from 0 to 60, with higher scores indicating more repetitive and restricted behaviours.

Participants also reported their anxiety and depressive symptoms with Hospital Anxiety and Depression Score (HADS)[6]. The HADS consists of 2 subscales for anxiety and depression, each including 7 items. Each item is rated with a 4-likert scale ranging from 0-3, adding up to a total score ranging from 0-21 for each subscale. Higher scores indicate more anxiety or depressive symptoms.

#### Data Analysis

##### *Task-based BOLD Response*

Subject-level whole-brain BOLD response to faces was characterised with general linear modelling implemented with FEAT toolbox of FSL v6.0.7.14. The regression model consisted of each stimulus type convolved with the double gamma hemodynamic response function. Contrast maps of parameter estimate were obtained from linear contrasts of interest: faces > shapes. Group-level general linear modelling with randomise permutation was performed using subject-level contrast maps as the dependent variable. The regression model consisted of the overall effect of faces (regardless of group or drug, using a voxel-wise family-wise-error (FWE) rate of one-tailed $p < 0.025$), main effect of group (including two contrasts of interest: autistic > non-autistic and autistic < non-autistic, using a voxel-wise FWE rate of one-tailed $p < 0.025$), age, gender, and mean FD (to control for residual motion effects) using FSL v6.0.7.14 randomise[7]. To examine drug and group × drug interaction effect, we subtracted the contrast maps under placebo from the contrast maps under psilocybin and then performed group-level general linear modelling with randomise permutation. The regression model consisted of the overall effect of drug (including two contrasts of interest: average > 0 and average < 0, each using a voxel-wise FWE rate of one-tailed $p < 0.025$), main effect of group, age, sex, and mean FD using FSL v6.0.7.14 randomise.

##### *Association with autistic traits*

When baseline difference or differential responses to psilocybin in a feature was found in autistic and non-autistic participants, we also examined the association of that feature or changes in that feature (5 mg psilocybin - placebo) with autistic traits and co-occurring mental health symptoms. We performed principal component analysis (PCA) on baseline scores of AASP, SRS-2, RBQ-2A, HADS-A, HADS-D from all the participants to obtain the first principal component (PC1), which explains the most variance in autistic core traits and associated mental health symptoms. PC1 scores of each participant were computed. We used multiple linear regression to examine the association between PC1 scores and ICN features.

#### fMRIprep preprocessing pipeline

Results included in this manuscript come from preprocessing performed using fMRIPrep 23.0.1 (Esteban et al. (2019); Esteban et al. (2018); RRID:SCR_016216), which is based on Nipype 1.8.5 (K. Gorgolewski et al. (2011); K. J. Gorgolewski et al. (2018); RRID:SCR_002502).

##### Anatomical data preprocessing

A total of 1 T1-weighted (T1w) images were found within the input BIDS dataset.The T1-weighted (T1w) image was corrected for intensity non-uniformity (INU) with N4BiasFieldCorrection (Tustison et al. 2010), distributed with ANTs 2.3.3 (Avants et al. 2008, RRID:SCR_004757), and used as T1w-reference throughout the workflow. The T1w-reference was then skull-stripped with a Nipype implementation of the antsBrainExtraction.sh workflow (from ANTs), using OASIS30ANTs as target template. Brain tissue segmentation of cerebrospinal fluid (CSF), white-matter (WM) and gray-matter (GM) was performed on the brain-extracted T1w using fast (FSL 6.0.5.1:57b01774, RRID:SCR_002823, Zhang, Brady, and Smith 2001). Brain surfaces were reconstructed using recon-all (FreeSurfer 7.3.2, RRID:SCR_001847, Dale, Fischl, and Sereno 1999), and the brain mask estimated previously was refined with a custom variation of the method to reconcile ANTs-derived and FreeSurfer-derived segmentations of the cortical gray-matter of Mindboggle (RRID:SCR_002438, Klein et al. 2017). Volume-based spatial normalization to two standard spaces (MNI152NLin6Asym, MNI152NLin2009cAsym) was performed through nonlinear registration with antsRegistration (ANTs 2.3.3), using brain-extracted versions of both T1w reference and the T1w template. The following templates were were selected for spatial normalization and accessed with TemplateFlow (23.0.0, Ciric et al. 2022): FSL's MNI ICBM 152 non-linear 6th Generation Asymmetric Average Brain Stereotaxic Registration Model [Evans et al. (2012), RRID:SCR_002823; TemplateFlow ID: MNI152NLin6Asym], ICBM 152 Nonlinear Asymmetrical template version 2009c [Fonov et al. (2009), RRID:SCR_008796; TemplateFlow ID: MNI152NLin2009cAsym].

##### Functional data preprocessing

For each of the 1 BOLD runs found per subject (across all tasks and sessions), the following preprocessing was performed. First, a reference volume and its skull-stripped version were generated using a custom methodology of fMRIPrep. Head-motion parameters with respect to the BOLD reference (transformation matrices, and six corresponding rotation and translation parameters) are estimated before any spatiotemporal filtering using mcflirt (FSL 6.0.5.1:57b01774, Jenkinson et al. 2002). BOLD runs were slice-time corrected to 0.462s (0.5 of slice acquisition range 0s-0.923s) using 3dTshift from AFNI (Cox and Hyde 1997, RRID:SCR_005927). The BOLD time-series (including slice-timing correction when applied) were resampled onto their original, native space by applying the transforms to correct for head-motion. These resampled BOLD time-series will be referred to as preprocessed BOLD in original space, or just preprocessed BOLD. The BOLD reference was then co-registered to the T1w reference using bbregister (FreeSurfer) which implements boundary-based registration (Greve and Fischl 2009). Co-registration was configured with twelve degrees of freedomto account for distortions remaining in the BOLD reference. Several confounding time-series were calculated based on the preprocessed BOLD: framewise displacement (FD), DVARS and three region-wise global signals. FD was computed using two formulations following Power (absolute sum of relative motions, Power et al. (2014)) and Jenkinson (relative root mean square displacement between affines, Jenkinson et al. (2002)). FD and DVARS are calculated for each functional run, both using their implementations in Nipype (following the definitions by Power et al. 2014). The three global signals are extracted within the CSF, the WM, and the whole-brain masks. Additionally, a set of physiological regressors were extracted to allow for component-based noise correction (CompCor, Behzadi et al. 2007). Principal components are estimated after high-pass filtering the preprocessed BOLD time-series (using a discrete cosine filter with 128s cut-off) for the two CompCor variants: temporal (tCompCor) and anatomical (aCompCor). tCompCor components are then calculated from the top 2% variable voxels within the brain mask. For aCompCor, three probabilistic masks (CSF, WM and combined CSF+WM) are generated in anatomical space. The implementation differs from that of Behzadi et al. in that instead of eroding the masks by 2 pixels on BOLD space, a mask of pixels that likely contain a volume fraction of GM is subtracted from the aCompCor masks. This mask is obtained by dilating a GM mask extracted from the FreeSurfer's aseg segmentation, and it ensures components are not extracted from voxels containing a minimal fraction of GM. Finally, these masks are resampled into BOLD space and binarized by thresholding at 0.99 (as in the original implementation). Components are also calculated separately within the WM and CSF masks. For each CompCor decomposition, the k components with the largest singular values are retained, such that the retained components' time series are sufficient to explain 50 percent of variance across the nuisance mask (CSF, WM, combined, or temporal). The remaining components are dropped from consideration. The head-motion estimates calculated in the correction step were also placed within the corresponding confounds file. The confound time series derived from head motion estimates and global signals were expanded with the inclusion of temporal derivatives and quadratic terms for each (Satterthwaite et al. 2013). Frames that exceeded a threshold of 0.5 mm FD or 1.5 standardized DVARS were annotated as motion outliers. Additional nuisance timeseries are calculated by means of principal components analysis of the signal found within a thin band (crown) of voxels around the edge of the brain, as proposed by (Patriat, Reynolds, and Birn 2017). The BOLD time-series were resampled into several standard spaces, correspondingly generating the following spatially-normalized, preprocessed BOLD runs: MNI152NLin6Asym, MNI152NLin2009cAsym. First, a reference volume and its skull-stripped version were generated using a custom methodology of fMRIPrep. Automatic removal of motion artifacts using independent component analysis (ICA-AROMA, Pruim et al. 2015) was performed on the preprocessed BOLD on MNI space time-series after removal of non-steady state volumes and spatial smoothing with an isotropic, Gaussian kernel of 6mm FWHM (full-width half-maximum). Corresponding “non-aggresively” denoised runs were produced after such smoothing. Additionally, the “aggressive” noise-regressors were collected and placed in the corresponding confounds file. All resamplings can be performed with a single interpolation step by composing all the pertinent transformations (i.e. head-motion transform matrices, susceptibility distortion correction when available, and co-registrations to anatomical and output spaces). Gridded (volumetric) resamplings were performed using antsApplyTransforms (ANTs), configured with Lanczos interpolation to minimize the smoothing effects of other kernels (Lanczos 1964). Non-gridded (surface) resamplings were performed using mri_vol2surf (FreeSurfer).

Many internal operations of fMRIPrep use Nilearn 0.9.1 (Abraham et al. 2014, RRID:SCR_001362), mostly within the functional processing workflow. For more details of the pipeline, see the section corresponding to workflows in fMRIPrep’s documentation.

#### Copyright Waiver

The above boilerplate text was automatically generated by fMRIPrep with the express intention that users should copy and paste this text into their manuscripts unchanged. It is released under the CC0 license.

Brain Models." Nature Methods 19: 1568–71. https://doi.org/10.1038/s41592-022-01681-2.

Cox, Robert W., and James S. Hyde. 1997. "Software Tools for Analysis and Visualization of fMRI Data." NMR in Biomedicine 10 (4-5): 171–78. https://doi.org/10.1002/(SICI)1099-1492(199706/08)10:4/5<171::AID-NBM453>3.0.CO;2-L.

Dale, Anders M., Bruce Fischl, and Martin I. Sereno. 1999. "Cortical Surface-Based Analysis: I. Segmentation and Surface Reconstruction." NeuroImage 9 (2): 179–94. https://doi.org/10.1006/nimg.1998.0395.

Esteban, Oscar, Ross Blair, Christopher J. Markiewicz, Shoshana L. Berleant, Craig Moodie, Feilong Ma, Ayse Ilkay Isik, et al. 2018. "fMRIPrep 23.0.1." Software. https://doi.org/10.5281/zenodo.852659.

Esteban, Oscar, Christopher Markiewicz, Ross W Blair, Craig Moodie, Ayse Ilkay Isik, Asier Erramuzpe Aliaga, James Kent, et al. 2019. "fMRIPrep: A Robust Preprocessing Pipeline for Functional MRI." Nature Methods 16: 111–16. https://doi.org/10.1038/s41592-018-0235-4.

Evans, AC, AL Janke, DL Collins, and S Baillet. 2012. "Brain Templates and Atlases." NeuroImage 62 (2): 911–22. https://doi.org/10.1016/j.neuroimage.2012.01.024.

Fonov, VS, AC Evans, RC McKinstry, CR Almli, and DL Collins. 2009. "Unbiased Nonlinear Average Age-Appropriate Brain Templates from Birth to Adulthood." NeuroImage 47, Supplement 1: S102. https://doi.org/10.1016/S1053-8119(09)70884-5.

Gorgolewski, K., C. D. Burns, C. Madison, D. Clark, Y. O. Halchenko, M. L. Waskom, and S. Ghosh. 2011. "Nipype: A Flexible, Lightweight and Extensible Neuroimaging Data Processing Framework in Python." Frontiers in Neuroinformatics 5: 13. https://doi.org/10.3389/fninf.2011.00013.

Gorgolewski, Krzysztof J., Oscar Esteban, Christopher J. Markiewicz, Erik Ziegler, David Gage Ellis, Michael Philipp Notter, Dorota Jarecka, et al. 2018. "Nipype." Software. https://doi.org/10.5281/zenodo.596855.

Greve, Douglas N, and Bruce Fischl. 2009. "Accurate and Robust Brain Image Alignment Using Boundary-Based Registration." NeuroImage 48 (1): 63–72. https://doi.org/10.1016/j.neuroimage.2009.06.060.

Jenkinson, Mark, Peter Bannister, Michael Brady, and Stephen Smith. 2002. "Improved Optimization for the Robust and Accurate Linear Registration and Motion Correction of Brain Images." NeuroImage 17 (2): 825–41. https://doi.org/10.1006/nimg.2002.1132.

Klein, Arno, Satrajit S. Ghosh, Forrest S. Bao, Joachim Giard, Yrjö Häme, Eliezer Stavsky, Noah Lee, et al. 2017. "Mindboggling Morphometry of Human Brains." PLOS Computational Biology 13 (2): e1005350. https://doi.org/10.1371/journal.pcbi.1005350.

Lanczos, C. 1964. “Evaluation of Noisy Data.” Journal of the Society for Industrial and Applied Mathematics Series B Numerical Analysis 1 (1): 76–85. https://doi.org/10.1137/0701007.

Patriat, Rémi, Richard C. Reynolds, and Rasmus M. Birn. 2017. “An Improved Model of Motion-Related Signal Changes in fMRI.” NeuroImage 144, Part A (January): 74–82. https://doi.org/10.1016/j.neuroimage.2016.08.051.

Power, Jonathan D., Anish Mitra, Timothy O. Laumann, Abraham Z. Snyder, Bradley L. Schlaggar, and Steven E. Petersen. 2014. “Methods to Detect, Characterize, and Remove Motion Artifact in Resting State fMRI.” NeuroImage 84 (Supplement C): 320–41. https://doi.org/10.1016/j.neuroimage.2013.08.048.

Pruim, Raimon H. R., Maarten Mennes, Daan van Rooij, Alberto Llera, Jan K. Buitelaar, and Christian F. Beckmann. 2015. “ICA-AROMA: A Robust ICA-Based Strategy for Removing Motion Artifacts from fMRI Data.” NeuroImage 112 (Supplement C): 267–77. https://doi.org/10.1016/j.neuroimage.2015.02.064.

Satterthwaite, Theodore D., Mark A. Elliott, Raphael T. Gerraty, Kosha Ruparel, James Loughead, Monica E. Calkins, Simon B. Eickhoff, et al. 2013. “An improved framework for confound regression and filtering for control of motion artifact in the preprocessing of resting-state functional connectivity data.” NeuroImage 64 (1): 240–56. https://doi.org/10.1016/j.neuroimage.2012.08.052.

Tustison, N. J., B. B. Avants, P. A. Cook, Y. Zheng, A. Egan, P. A. Yushkevich, and J. C. Gee. 2010. “N4itk: Improved N3 Bias Correction.” IEEE Transactions on Medical Imaging 29 (6): 1310–20. https://doi.org/10.1109/TMI.2010.2046908.

Zhang, Y., M. Brady, and S. Smith. 2001. “Segmentation of Brain MR Images Through a Hidden Markov Random Field Model and the Expectation-Maximization Algorithm.” IEEE Transactions on Medical Imaging 20 (1): 45–57. https://doi.org/10.1109/42.906424
